# The potassium channel KCNJ13 is essential for smooth muscle cytoskeletal organization during mouse tracheal tubulogenesis

**DOI:** 10.1101/320119

**Authors:** Wenguang Yin, Hyun-Taek Kim, ShengPeng Wang, Felix Gunawan, Lei Wang, Keishi Kishimoto, Hua Zhong, Dany Roman, Jens Preussner, Stefan Guenther, Viola Graef, Carmen Buettner, Beate Grohmann, Mario Looso, Mitsuru Morimoto, Graeme Mardon, Stefan Offermanns, Didier Y.R. Stainier

## Abstract

Tubulogenesis is essential for the formation and function of internal organs. One such organ is the trachea, which allows gas exchange between the external environment and the lungs. However, the cellular and molecular mechanisms underlying tracheal tube development remain poorly understood. Here, we show that the potassium channel KCNJ13 is a critical modulator of tracheal tubulogenesis. We identify *Kcnj13* in an ethylnitrosourea forward genetic screen for regulators of mouse respiratory organ development. *Kcnj13* mutants exhibit a shorter trachea as well as defective smooth muscle (SM) cell alignment and polarity. KCNJ13 is essential to maintain ion homeostasis in tracheal SM cells, which is required for actin polymerization. This process appears to be mediated, at least in part, through activation of the actin regulator AKT, as pharmacological increase of AKT phosphorylation ameliorates the *Kcnj13* mutant trachea phenotypes. These results provide insights into the role of ion homeostasis in cytoskeletal organization during tubulogenesis.

## Introduction

The trachea consists of endoderm-derived epithelium surrounded by mesoderm-derived cartilage, connective tissue and smooth muscle (SM)^1^. The SM provides the elasticity necessary to control tracheal contraction, whereas the cartilage provides tissue rigidity to prevent airway collapse^2^. In humans, tracheal tube formation defects have been reported to cause tracheomalacia, which is characterized by deficiency of the supporting cartilage and may lead to airway collapse, respiratory distress, and death^3^. Studies on the cellular and molecular mechanisms underlying tracheal tubulogenesis have mostly focused on the role of epithelial cells^4,5^ as well as the complex signaling between the epithelium and mesenchyme^6,7^, but how SM cells regulate this process remains unknown.

Another poorly studied aspect of tubulogenesis is the potential role of ion channels and their mediated ion homeostasis. Recent data indicate that potassium channels play important roles in the behavior of non-excitable cells, including in tissue patterning^8,9^. In this context, cytoskeletal organization, including the modulation of actin dynamics, is essential to maintain cell shape and alignment. Amongst a variety of actin-associated factors, the serine/threonine kinase AKT has emerged as one important regulator of actin organization^10,11^. AKT phosphorylates Girdin, an actin-binding protein that regulates F-actin levels, and phosphorylated Girdin accumulates at the leading edge of migrating cells^11^. These data provide a direct link between AKT activity and actin polymerization. However, how AKT activity is regulated is poorly understood.

Here, starting with a forward genetic approach in mouse, we reveal a role for a specific potassium channel, and ion homeostasis, during tracheal tubulogenesis, at least in part via the control of actin dynamics and cytoskeletal organization.

## Results

### *Kcnj13^T38C/T38C^* mice exhibit tracheal defects

We sought to identify regulators of mouse respiratory organ formation by conducting a large-scale forward genetic screen using ethylnitrosourea (ENU) mutagenesis. Mutagenized C57BL/6J mice were bred to uncover recessive phenotypes using a two generation backcross breeding scheme (Supplementary Fig. 1a). Early postnatal tracheas were examined using alcian blue to label cartilage, and alpha smooth muscle actin (αSMA) immunostaining in whole-mount preparations (Fig. 1a). We screened 473 G1 animals and recovered 5 mutants with tracheal tube formation defects, one of which displayed cyanosis (Fig. 1b) and neonatal respiratory distress (Fig. 1c and Supplementary Movie 1), and died within 24 hours of birth. These mutant animals were born in the expected Mendelian ratio with a shortened (Fig. 1d, e) and collapsed (Fig. 1f, g) trachea, and fractured cartilage rings instead of the intact ventrolateral cartilage rings seen in wild-type (WT) siblings (Fig. 1d, f). In addition, disorganized SM, characterized by the narrowing of SM stripes, was observed in the mutants (Fig. 1h). To examine the lungs, we performed hematoxylin and eosin staining of tissue sections. Mutants exhibited reduced air space and thickened walls in distal regions compared to WT (Fig. 1i, j). These data indicate that the mutants die from compromised respiratory function due to defective formation of the tracheal tube and respiratory air sacs.

**Figure 1.**
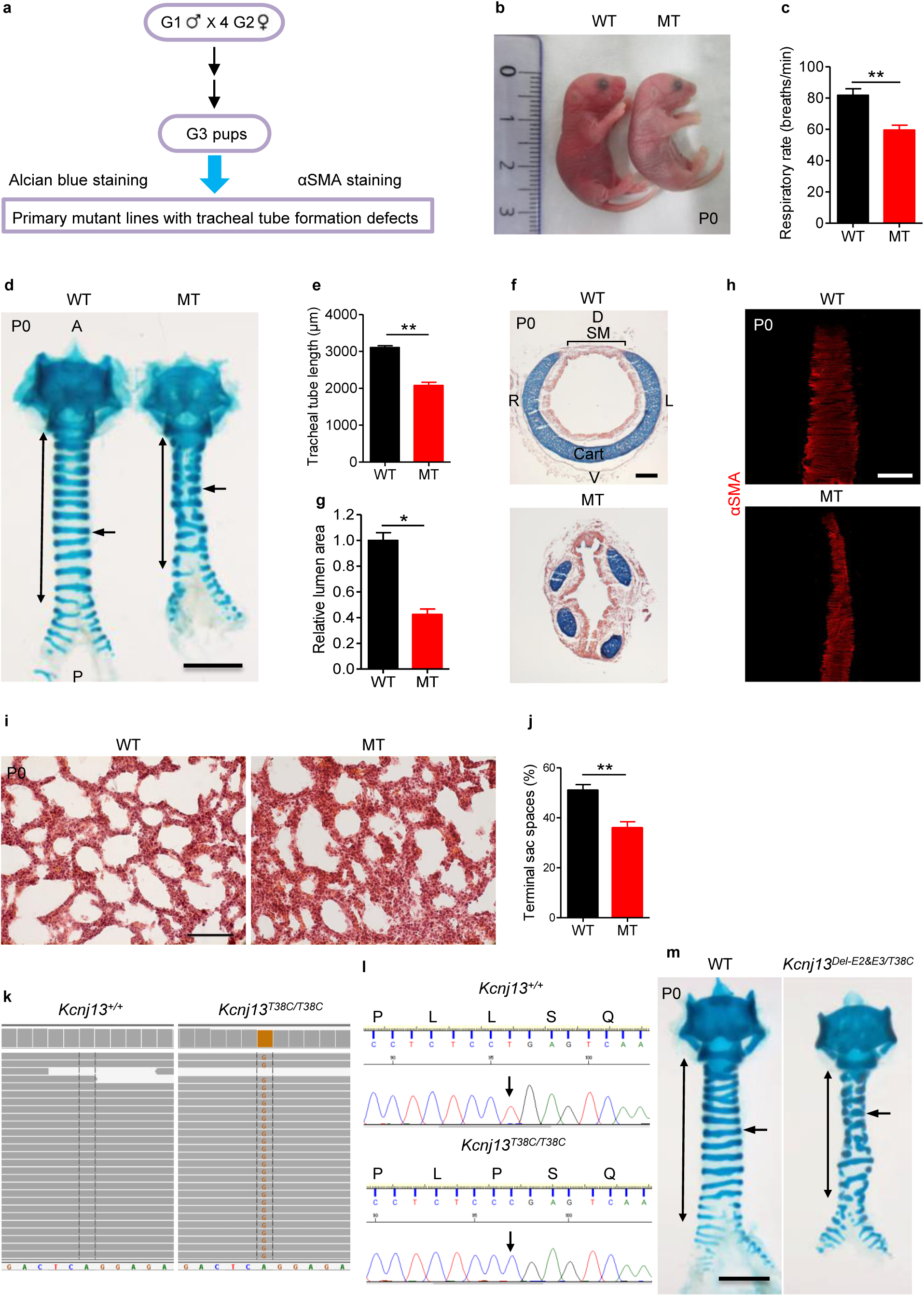
*Kcnj13^T38C/T38C^* mice exhibit tracheal tube formation defects. (**a**) Workflow for ENU screen in mouse to identify mutants with respiratory organ formation defects. (**b**) Representative gross morphology of P0 WT (n=12) and mutants (n=12). (**c**) Quantification of P0 WT (n=12) and mutant (n=12) respiratory rates. (**d**) Representative image of ventral view of wholemount tracheas stained with alcian blue from P0 WT (n=35) and mutants (n=35). Double-sided arrows indicate tracheal tube length. Arrows point to tracheal cartilage rings. (**e**) Quantification of P0 WT (10) and mutant (10) tracheal tube length. (**f**) Representative images of transverse sections of tracheas stained with alcian blue and nuclear fast red from P0 WT (n=9) and mutants (n=9). (**g**) Quantification of P0 WT (n=9) and mutant (9) tracheal lumen area. (**h**) Representative images of dorsal views of wholemount tracheas stained for αSMA (red) from P0 WT (n=10) and mutants (n=10). (**i**) Representative images of lung tissue sections stained with hematoxylin and eosin from P0 WT (n=8) and mutants (n=8). (**j**) Quantification of P0 WT (n=8) and mutant (8) terminal airspaces in the lungs. (**k**) Whole-exome sequencing of P0 WT (n=2) and mutant (n=2) genomic DNA. Green indicates the WT nucleotide A. Light brown indicates the mutant nucleotide G. (**l**) Sequence of WT and mutant genomic DNA around the lesion. DNA sequence fluorogram shows CTG for leucine in WT (n=119) (upper panel) and CCG for proline in mutants (n=105) (lower panel). Arrows point to the mutation site. (**m**) Representative images of ventral views of wholemount tracheas stained with alcian blue from P0 WT (n=17) and *Kcnj13^Del-E2&E3/T38C^* double heterozygous animals (n=17). Double-sided arrows indicate tracheal tube length. Arrows point to tracheal cartilage rings. Scale bars: 1000 μm (**d, m**), 500 μm (**h**), 100 μm (**e, i**). \**P* < 0.05; \*\**P* < 0.01; Unpaired Student’s *t*-test, mean ± s.d. WT, Wild-type; MT, Mutant; A, Anterior; P, Posterior; D, Dorsal; V, Ventral; L, Left; R, Right; Cart, Cartilage; SM, Smooth muscle.

To identify the phenotype-causing mutation, we conducted whole-exome sequencing of G4 genomic DNA samples (Supplementary Fig. 1b), and identified *Kcnj13*, which encodes a member of the inwardly rectifying potassium channel family, as a candidate gene (Fig. 1k). The identified allele carries a mutation that causes a leucine-to-proline substitution at the highly conserved 13^th^ residue (c.38T>C (p.Leu13Pro)). Next, we carried out genetic linkage analysis by genotyping 105 G4 and G5 mutant animals, and found complete linkage between the tracheal phenotype and the *Kcnj13^T38C/T38C^* allele (Fig. 1l). We then performed a complementation test by crossing mice carrying the ENU-induced *Kcnj13* allele (*Kcnj13^T38C/+^*) with mice carrying a *Kcnj13* deletion allele (Supplementary Fig. 2a-d), and found that complementation did not occur in the *Kcnj13^Del-E2&E3/T38C^* double heterozygous animals (Fig. 1m), indicating that loss of KCNJ13 function is likely responsible for the observed tracheal phenotypes. To further test the role of *Kcnj13* in tracheal development, we analyzed tracheal formation in *Kcnj13^Del-E2&E3^* mice. *Kcnj13^Del-E2&E3^* mice exhibited a shortened trachea with fractured cartilage rings (Supplementary Fig. 3a, b) and disorganized SM (Supplementary Fig. 3c, d), and died within 24 hours of birth, similar to *Kcnj13^T38C/T38C^* animals. We then examined the spatial and temporal expression pattern of *Kcnj13* in the developing mouse trachea and lungs. *Kcnj13* mRNA was expressed at low levels in E11.5-E13.5 tracheas (Supplementary Fig. 4a and Supplementary Table 1). At E13.5 and E14.5, KCNJ13 protein was clearly detected in tracheal SM (Fig. 2a), but not in SOX9^+^ mesenchymal cells (Fig. 2b). After E15.5, KCNJ13 was also detected in a subset of tracheal epithelial cells (Fig. 2c). In the lungs, *Kcnj13* mRNA was detected in epithelial cells as early as E16.5 (Supplementary Fig. 4b). KCNJ13 was expressed in epithelial cells of bronchioles (Fig. 2d, e) and alveolar type II cells (Fig. 2f), similar to the expression of a *Kcnj13* BAC reporter^12^. Interestingly, KCNJ13 protein was still detectable at WT levels in *Kcnj13^T38C/T38C^* tracheas (Supplementary Fig. 4c), indicating that the c.38T>C mutation has no effect on protein stability. Altogether, these results indicate that loss of *Kcnj13* function causes severe tracheal defects.

**Figure 2.**
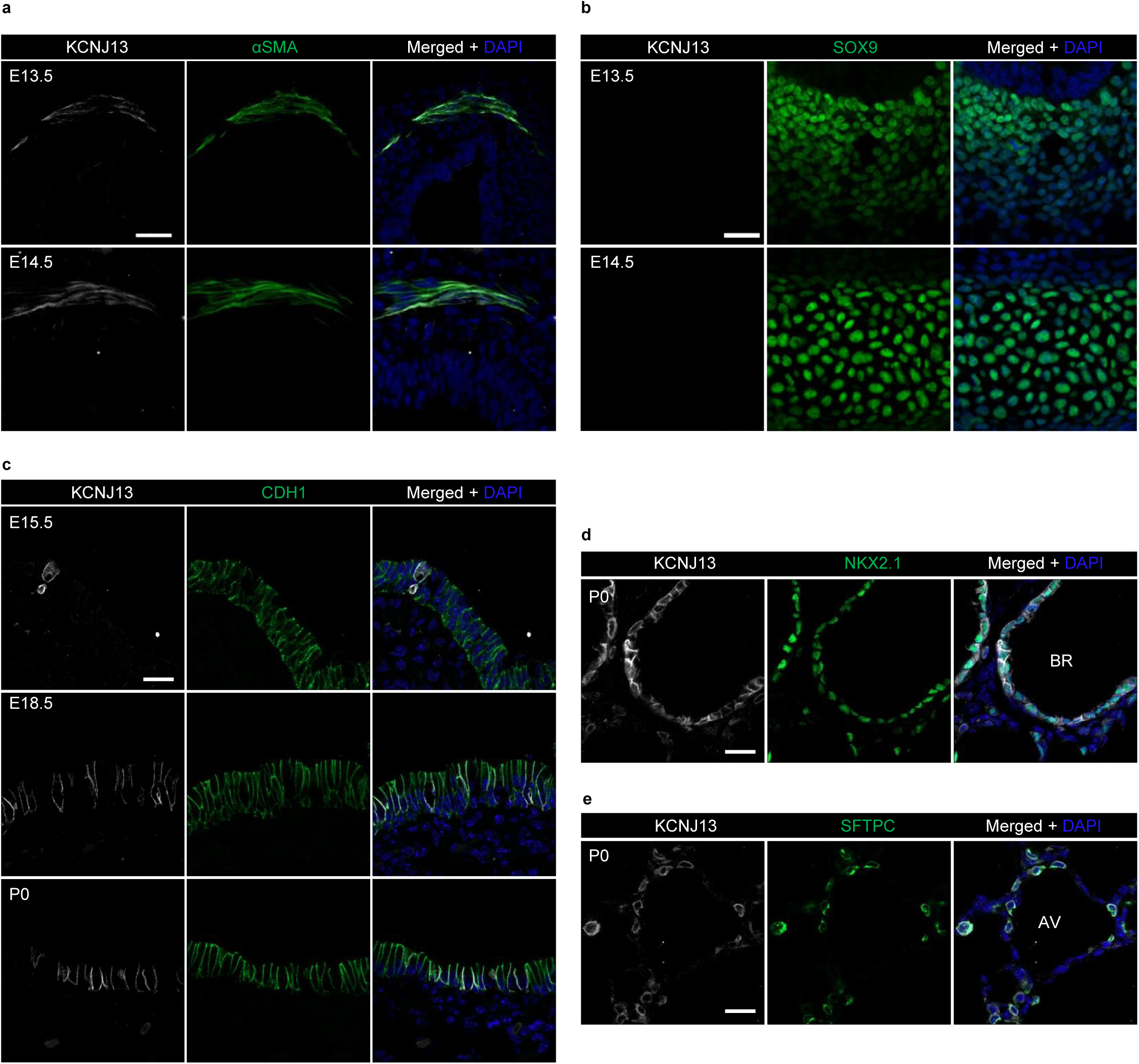
*Kcnj13* is expressed in SM and epithelial cells in the trachea, as well as in epithelial cells in the lungs. (**a**) Immunostaining for KCNJ13 (red), αSMA (green) and DAPI staining (blue) of transverse sections of E13.5 (n=8) and E14.5 (n=8) WT tracheas. (**b**) Immunostaining for KCNJ13 (red), SOX9 (green) and DAPI staining (blue) of transverse sections of E13.5 (n=8) and E14.5 (n=8) WT tracheas. (**c**) Immunostaining for KCNJ13 (red), CDH1 (green) and DAPI staining (blue) of transverse sections of E15.5 (n=8), E18.5 (n=8) and P0 (n=8) WT tracheas. (**d**) Immunostaining for KCNJ13 (red), NKX2.1 (green) and DAPI staining (blue) of P0 WT lung tissue sections (n=7). (**e**) Immunostaining for KCNJ13 (red), SFTPC (green) and DAPI staining (blue) of P0 WT lung tissue sections (n=7). Scale bars: 20 μm. BR, Bronchioles; AV, Alveoli.

### *Kcnj13^T38C/T38C^* mice exhibit defects in tracheal elongation

To examine the formation of the trachea in detail, we performed a systematic analysis of tracheal tube development. Tracheal tube length did not significantly differ between *Kcnj13^T38C/T38C^* mice and their WT siblings from E11.5 to E13.5 (Fig. 3a, b). However, starting at E14.5, we observed that *Kcnj13^T38C/T38C^* tracheas were shorter than WT (Fig. 3a, b), indicating that impaired tracheal tube elongation occurs after SM differentiation, which starts at E11.5^2^. Since altered SM morphogenesis can affect tracheal elongation^13^, we analyzed tracheal SM development. SM cells which are positioned dorsally in the trachea, displayed no obvious differences between WT and *Kcnj13^T38C/T38C^* animals from E11.5 to E12.5 (Fig. 3c, d). Disorganized SM stripes of decreased area were first observed in E13.5 *Kcnj13^T38C/T38C^* tracheas and became more noticeable starting at E14.5 (Fig. 3c, d). Another important process during trachea formation is the appearance of cartilage from the condensation of mesenchymal cells into chondrogenic nodules^6^. At E13.5, a clear pattern of condensed SOX9^+^ mesenchymal cells resembling cartilaginous rings was readily distinguished in WT, whereas such condensations were seldom detected in *Kcnj13^T38C/T38C^* tracheas (Fig. 3e, f). This effect on SM organization and mesenchymal condensation appeared specific, as we did not observe significant differences between WT and mutant animals in the proliferation of SM cells (Supplementary Fig. 5a, b) or SOX9^+^ mesenchymal cells (Supplementary Fig. 5c, d) or apoptosis (Supplementary Fig. 5e). Next, we examined the expression levels of *Sox9*, which encodes a key regulator of chondrogenic nodule formation^14^. *Kcnj13^T38C/T38C^* tracheas exhibited no significant difference in *Sox9* mRNA levels compared to WT (Supplementary Fig. 5f and Supplementary Table 2), indicating that *Kcnj13* participates in cartilage formation through a *Sox9* independent pathway. Potassium channels have been reported to modulate cell differentiation^15^. We observed reduced differentiation of acetylated alpha-tubulin^+^ multiciliated cells (Supplementary Fig. 5g, h) but a WT-like distribution of CC10^+^ club cells (Supplementary Fig. 5g, i) and KRT5^+^ basal cells (Supplementary Fig. 5g, j) in the mutant tracheas. In addition, *Kcnj13^T38C/T38C^* tracheas exhibited WT-like organization of the tracheal epithelium (Supplementary Fig. 5k). Collectively, these results suggest that SM organization and mesenchymal condensation are essential for tracheal tube formation.

**Figure 3.**
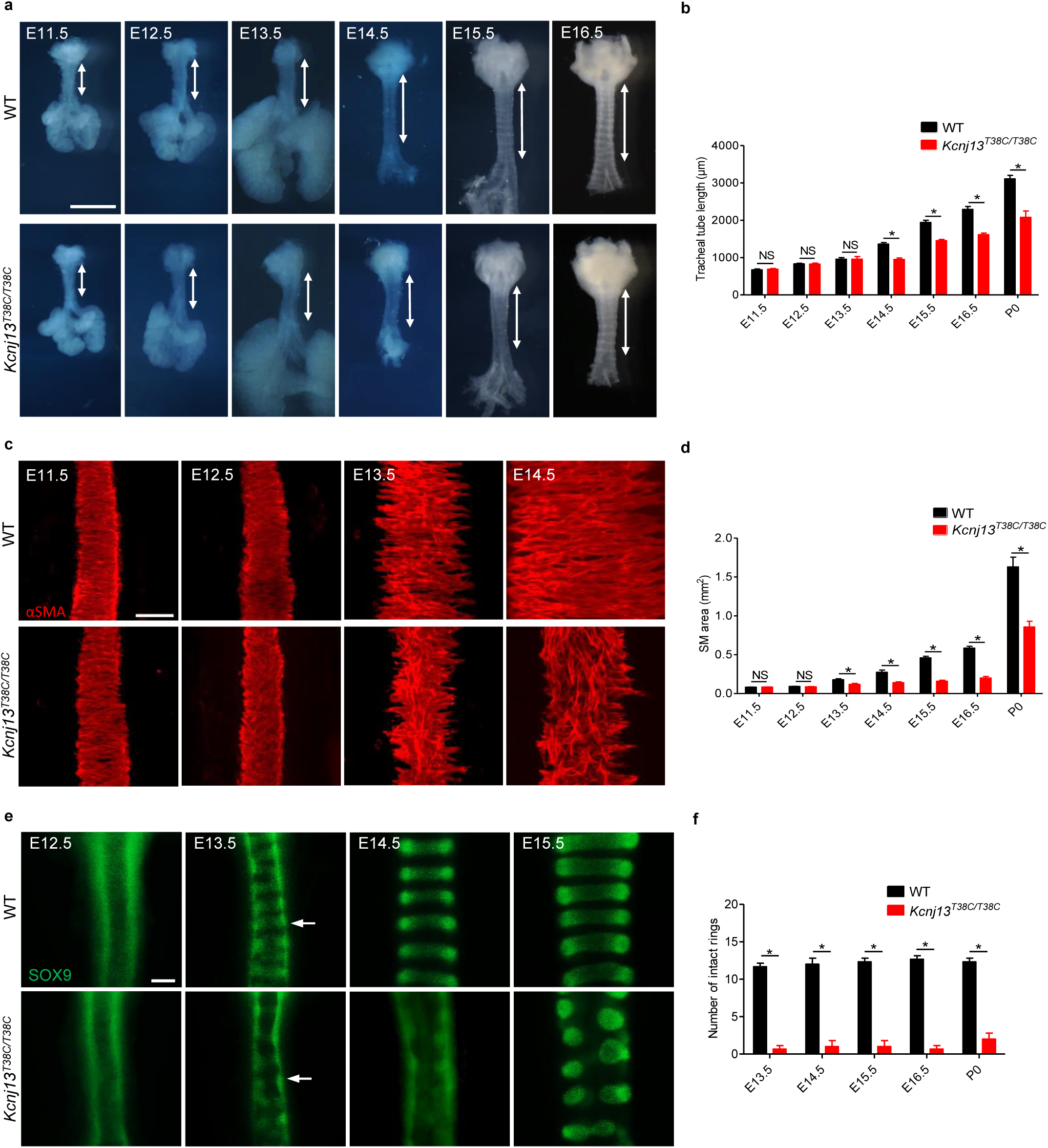
*Kcnj13^T38C/T38C^* mice exhibit defects in tracheal elongation, SM organization, and mesenchymal condensation. (**a**) Representative images of ventral views of WT (n=8) and *Kcnj13^T38C/T38C^* (n=8) tracheas at several embryonic stages. Double-sided arrows indicate tracheal tube length. (**b**) Quantification of WT (n=8) and *Kcnj13^T38C/T38C^* (n=8) tracheal tube length. (**c**) Immunostaining for αSMA (red) in dorsal views of WT (n=8) and *Kcnj13^T38C/T38C^* (n=8) tracheas at several embryonic stages. (**d**) Quantification of WT (n=8) and *Kcnj13^T38C/T38C^* (n=8) SM area. (**e**) Immunostaining for SOX9 (green) in ventral views of WT (n=8) and *Kcnj13^T38C/T38C^* (n=8) tracheas at several embryonic stages. Arrows point to tracheal mesenchymal condensations. (**f**) Quantification of intact tracheal mesenchymal condensations in WT (n=8) and *Kcnj13^T38C/T38C^* (n=8) animals. Scale bars: 1000 μm (**a**), 100 μm (**c, e**). \**P* < 0.05; NS, not significant. Unpaired Student’s *t*-test, mean ± s.d.

### *Kcnj13* modulates SM cell alignment and polarity

To investigate the cellular mechanisms underlying *Kcnj13*-mediated tracheal tube formation, we analyzed SM cell alignment and polarity in WT and *Kcnj13^T38C/T38C^* tracheas. Tracheal SM cells differentiate and acquire radial cell polarity starting at E11.5^13^; they develop spindle shapes and become circumferentially aligned by E13.5 (Fig. 3c). WT tracheal SM cells were aligned in a direction perpendicular to that of tracheal elongation by E14.5 (Fig. 4a, b). In contrast, *Kcnj13^T38C/T38C^* SM cells displayed random alignment (Fig. 4a), which could be quantitatively assessed (Fig. 4b), as well as abnormally rounded nuclei (Fig. 4a, c). These mutant SM cells aligned into 7–8 layers compared to 3–4 layers in WT at E14.5 (Fig. 4d, e). To better understand the polarization of SM cells, we examined the localization of the Golgi apparatus relative to the cell nucleus, using the cis-Golgi matrix marker GM130, a widely used method to determine cell polarity in different cell types^16–18^. In WT SM cells, the GM130-labeled Golgi exhibited a ribbon-like morphology and localized preferentially by the long edges of the nucleus (Fig. 4f, g). In contrast, in *Kcnj13^T38C/T38C^* SM cells, the Golgi exhibited a more compact structure with random alignment (Fig. 4f, g). Since Wnt5a-Ror2 signaling modulates tracheal SM cell polarity^13^, we examined for changes in Wnt/planar cell polarity (PCP), a pathway known to control cell polarity^19^. *Kcnj13^T38C/T38C^* tracheas exhibited no significant differences in expression levels of Wnt/PCP genes (Supplementary Fig. 6 and Supplementary Table 3), indicating that *Kcnj13* signals via a Wnt/PCP independent pathway to direct SM cell polarity. In addition, SOX9^+^ mesenchymal cells displayed a WT-like morphology in *Kcnj13^T38C/T38C^* tracheas (Supplementary Fig. 7a, b). These findings indicate that KCNJ13 is required to coordinate SM cell polarity and generate tracheal tissue architecture.

**Figure 4.**
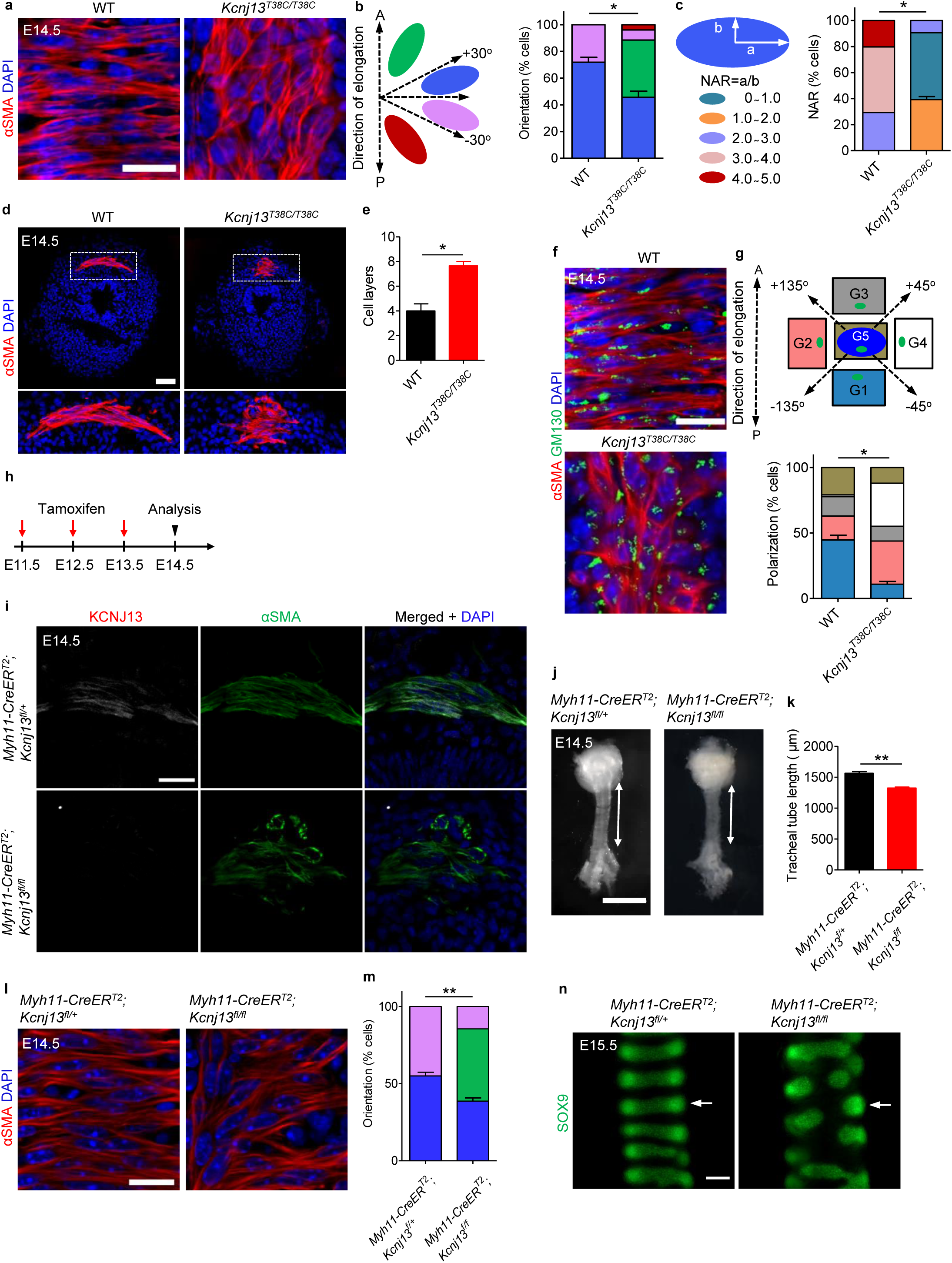
*Kcnj13* orchestrates SM cell alignment and polarity. (**a**) Immunostaining for αSMA (red) and DAPI staining (blue) in dorsal views of E14.5 WT (n=8) and *Kcnj13^T38C/T38C^* (n=8) tracheas. (**b**) Quantification of E14.5 WT (n=8) and *Kcnj13^T38C/T38C^* (n=8) tracheal SM cell orientation. (**c**) Quantification of E14.5 WT (n=8) and *Kcnj13^T38C/T38C^* (n=8) tracheal SM cell nuclear aspect ratio (NAR). (**d**) Immunostaining for αSMA (red) and DAPI staining (blue) of transverse sections of E14.5 WT (n=10) and *Kcnj13^T38C/T38C^* (n=10) tracheas. (**e**) Quantification of WT (n=10) and *Kcnj13^T38C/T38C^* (n=10) tracheal SM cell layers. (**f**) Immunostaining for αSMA (red) and GM130 (green) and DAPI staining (blue) in dorsal views of E14.5 WT (n=8) and *Kcnj13^T38C/T38C^* (n=8) tracheas. (**g**) Quantification of WT (n=8) and *Kcnj13^T38C/T38C^* (n=8) Golgi apparatus (green) position relative to the nucleus (blue). (**h**) Timeline for tamoxifen administration. (**i**) Immunostaining for KCNJ13 (red), αSMA (green) and DAPI staining (blue) of transverse sections of E14.5 control (n=6) and *Myh11****-****CreER^T2^;Kcnj13 ^flox/flox^* (n=6) tracheas. (**j**) Representative images of ventral views of E14.5 control (n=9) and *Myh11****-****CreER^T2^;Kcnj13 ^flox/flox^* (n=9) tracheas. Double-sided arrows indicate tracheal tube length. (**k**) Quantification of E14.5 control (n=9) and *Myh11****-****CreER^T2^;Kcnj13 ^flox/flox^* (n=9) tracheal tube length. (**l**) Immunostaining for αSMA (red) and DAPI staining (blue) in dorsal views of E14.5 control (n=9) and *Myh11***-***CreER^T2^;Kcnj13 ^flox/flox^* (n=9) tracheas. (**m**) Quantification of E14.5 control (n=9) and *Myh11***-***CreER^T2^;Kcnj13 ^flox/flox^* (n=9) tracheal SM cell orientation. (**n**) Immunostaining for SOX9 (green) in ventral views of E15.5 control (n=6) and *Myh11***-***CreER^T2^;Kcnj13 ^flox/flox^* (n=6) tracheas. Arrows point to tracheal mesenchymal condensations. Scale bars: 1000 μm (**j**), 100 μm (**n**), 50 μm (**d**), 20 μm (**a, f, i, l**). \**P* < 0.05; \*\**P* < 0.01; Unpaired Student’s *t*-test, mean ± s.d.

To investigate whether defects in condensation of SOX9^+^ mesenchymal cells might affect tracheal tube elongation, we deleted Sonic hedgehog (*Shh*) function in the tracheal epithelium using *Nkx2.1^Cre^;Shh^flox/flox^* mice^6^. *Shh* is essential for tracheal cartilage formation^20^ and epithelial deletion of *Shh* using *Nkx2.1^Cre^;Shh^flox/flox^* mice leads to mesenchymal cell condensation defects^6^. As reported, *Nkx2.1^Cre^;Shh^flox/flox^* tracheas displayed severe SOX9^+^ mesenchymal cell condensation defects compared to controls at E16.5 (Supplementary Fig. 8a, b), as observed in *Kcnj13^T38C/T38C^* tracheas. However, SM cell alignment (Supplementary Fig. 8c, d) or tracheal tube length (Supplementary Fig. 8e, f) did not appear to be affected in these animals, indicating that mesenchymal cell condensation is not required for tracheal tube elongation.

To test whether the SM cell defects are responsible for the tracheal tube elongation phenotype in *Kcnj13^T38C/T38C^* mice, we used the SM-specific Cre mouse line *Myh11***-***CreER^T2^*, which induces efficient recombination in airway SM cells^21^. We conditionally deleted *Kcnj13* in SM by intraperitoneal tamoxifen injection for 3 consecutive days (2 mg per day) from E11.5 to E13.5 (Fig. 4h). Mice with SM-specific *Kcnj13* deletion (Fig. 4i) exhibited shorter tracheas (Fig. 4j, k), altered SM cell alignment (Fig. 4l, m) and cartilage formation defects (Fig. 4n) compared to controls, phenotypes that were not observed when *Kcnj13* was deleted in epithelial cells (*Nkx2.1^Cre^;Kcnj13^flox/flox^*) (Supplementary Fig. 8g-o). Moreover, *Kcnj13^T38C/T38C^* mice exhibited no significant differences in mitotic spindle orientation (Supplementary Fig. 9a, b) or cell proliferation in the tracheal epithelium (Supplementary Fig. 9c, d), or key growth factor gene expression (Supplementary Fig. 9e and Supplementary Table 4), indicating that *Kcnj13* mutations do not affect cell growth in the epithelium. Altogether, these data indicate that KCNJ13 function is specifically required in SM, but not in epithelial cells, for tracheal tube elongation.

Both the trachea and esophagus are derived from the foregut and separate after E9.5^22,23^. Next, we sought to determine whether *Kcnj13* was also required for esophageal elongation. We measured P0 esophagi and found that *Kcnj13^T38C/T38C^* mice displayed shortened esophageal tubes compared to their WT siblings (Supplementary Fig. 10a, b). We also analyzed the expression pattern of KCNJ13 in the developing esophagus: KCNJ13 was weakly expressed in E12.5 esophageal SM cells and this expression level increased as the SM tissue developed (Supplementary Fig. 10c). Next, we examined esophageal SM morphology. Newly differentiated esophageal SM cells were not fully elongated or well organized at E11.5-E12.5 (Supplementary Fig. 10d, e). They developed spindle shapes and became circumferentially aligned by E13.5 (Supplementary Fig. 10d, e). We found that esophageal SM was disorganized in *Kcnj13^T38C/T38C^* mice (Supplementary Fig. 10f) with altered SM cell alignment (Supplementary Fig. 10g, h) and polarity (Supplementary Fig. 10i, j). These data indicate that KCNJ13-mediated SM cell alignment and polarity play a broader role in epithelial tubulogenesis.

### *Kcnj13* modulates actin organization in SM cells

Inactivation of Kir channels has been reported to cause depolarization of cell membranes^24^. To determine the molecular mechanisms underlying KCNJ13-regulated SM cell alignment and polarity, we examined SM cell membrane potential by using the voltage-sensitive fluorescent dye DiBAC_4_ (3)^25^. The fluorescence intensity appeared higher in *Kcnj13^T38C/T38C^* SM cells compared to WT (Fig. 5a, b), suggesting that mutant SM cell membranes are depolarized. In addition, after treatment with 50 μM VU590, a KCNJ13 inhibitor^26^, E14.5 WT tracheas also exhibited SM cell membrane depolarization compared to controls (Fig. 5c, d).

**Figure 5.**
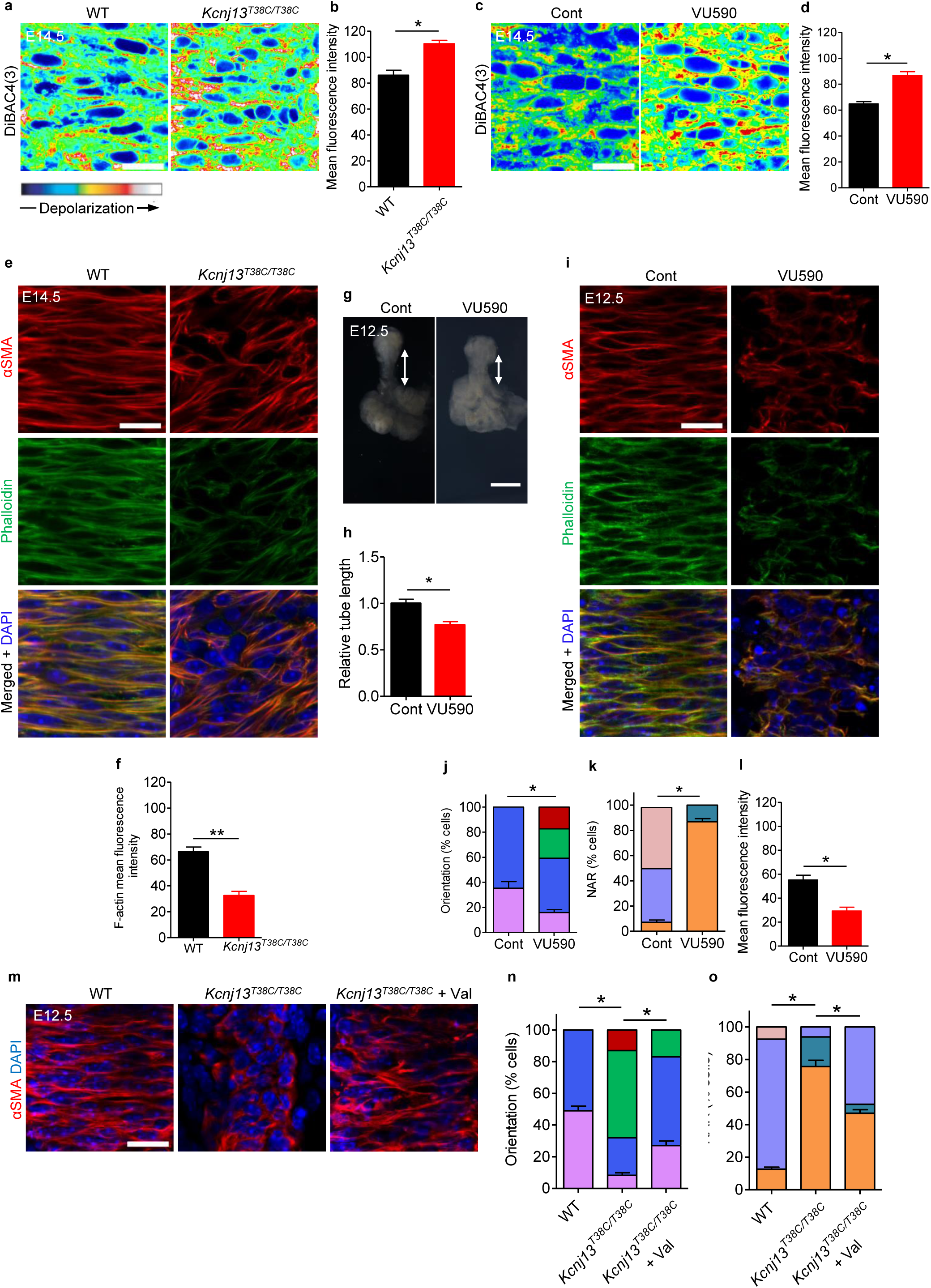
*Kcnj13* regulates membrane potential and actin organization in SM cells. (**a**) DiBAC4(3) fluorescence in dorsal views of E14.5 WT (n=9) and *Kcnj13^T38C/T38C^* (n=9) tracheal SM cells. (**b**) Quantification of mean DiBAC4(3) fluorescence intensity in E14.5 WT (n=9) and *Kcnj13^T38C/T38C^* (n=9) tracheal SM cells. (**c**) DiBAC4(3) fluorescence in dorsal views of E14.5 tracheal SM cells after DMSO (n=6) or 50 μM VU590 (n=6) treatment. (**d**) Quantification of mean DiBAC4(3) fluorescence intensity in E14.5 tracheal SM cells after DMSO (n=6) or 50 μM VU590 (n=6) treatment. (**e)** Immunostaining for αSMA (red) and phalloidin (green) and DAPI (blue) staining in dorsal views of E14.5 WT (n=8) and *Kcnj13^T38C/T38C^* (n=8) tracheas. (**f**) Quantification of mean phalloidin fluorescence intensity in E14.5 WT (n=8) and *Kcnj13^T38C/T38C^* (n=8) tracheal SM cells. (**g**) Representative images of ventral views of E12.5 tracheas after a 48 hour DMSO (n=6) or 50 μM VU590 (n=6) treatment. Double-sided arrows indicate tracheal tube length. (**h**) Quantification of relative tracheal tube length after a 48 hour DMSO (n=6) or 50 μM VU590 (n=6) treatment. (**i**) Immunostaining for αSMA (red) and phalloidin (green) and DAPI (blue) staining in dorsal views of E12.5 tracheas after 48 hour DMSO (n=9) or 50 μM VU590 (n=9) treatment. Quantification of E12.5 tracheal SM cell orientation (**j**), NAR (**k**), and mean phalloidin fluorescence intensity (**l**) after a 48 hour DMSO (n=9) or 50 μM VU590 (n=9) treatment. (**m**) Immunostaining for αSMA (red) and DAPI staining (blue) in dorsal views of E12.5 WT (n=24) and *Kcnj13^T38C/T38C^* (n=12) tracheas after a 48 hour DMSO treatment, and *Kcnj13^T38C/T38C^* tracheas (n=12) after a 48 hour 2 μM ionophore valinomycin treatment. Quantification of SM cell orientation (**n**) and NAR (**o**) of E12.5 WT (n=12) and *Kcnj13^T38C/T38C^* (n=12) tracheas after a 48 hour DMSO treatment, and *Kcnj13^T38C/T38C^* tracheas (n=12) after a 48 hour 2 μM ionophore valinomycin treatment. Scale bars: 500 μm (**e**), 20 μm (**a, c, g, i, m**). \**P* < 0.05; \*\**P* < 0.01; Unpaired Student’s *t*-test, mean ± s.d. Cont, Control; Val, Valinomycin.

Membrane depolarization has been reported to decrease the amount of actin filaments (F-actin)^27^ and thus affect mechanical support and cell shape^28^. Thus, we hypothesized that SM cell membrane depolarization might lead to actin depolymerization, and consequently to alteration of cell shape and alignment. We examined F-actin content by phalloidin staining, and observed that *Kcnj13^T38C/T38C^* tracheas exhibited decreased F-actin levels in SM cells compared to WT (Fig. 5e, f). In addition, an *ex vivo* 48 hour VU590 treatment led to a shortened trachea (Fig. 5g, h) with altered SM cell alignment (Fig. 5i, j) and shape (Fig. 5i, k), as well as decreased F-actin levels (Fig. 5i, l), similar to the phenotypes observed in *Kcnj13^T38C/T38C^* tracheas. These data indicate that KCNJ13-regulated membrane potential modulates actin organization in tracheal SM cells.

KCNJ13, reported to have unique pore properties^29^, has been shown to facilitate the efflux of intracellular potassium^30,31^. Thus, we hypothesized that accumulation of intracellular positive charges was responsible for the altered SM cell alignment and shape phenotypes in *Kcnj13^T38C/T38C^* tracheas. To test this hypothesis, we used valinomycin, a potassium ionophore reported to reduce intracellular potassium levels^32^, to deplete intracellular potassium in *Kcnj13^T38C/T38C^* tracheas in an *ex vivo* tracheal-lung organ culture system^33^. After 2 μM valinomycin treatment, *Kcnj13^T38C/T38C^* tracheas exhibited partially rescued SM cell alignment (Fig. 5m, n) and shape (Fig. 5m, o) phenotypes compared to DMSO-treated *Kcnj13^T38C/T38C^* tracheas. These data indicate that KCNJ13-mediated intracellular ion homeostasis is essential for tracheal SM cell alignment and shape.

Elevated extracellular potassium concentration has been reported to induce cell depolarization^34^ and increase intracellular potassium levels^32^. We hypothesized that an increase in extracellular potassium concentration might phenocopy the KCNJ13 inactivation-induced tracheal SM cell defects. We examined SM cell membrane potential in E14.5 tracheas after 40 mM KCl treatment. The intensity of fluorescence emitted from DiBAC4(3) was higher in tracheal SM cells after KCl treatment compared to controls (Supplementary Fig. 11a, b). Interestingly, after a 48 hour 40 mM KCl treatment, E12.5 tracheas exhibited a narrowing of SM stripes (Supplementary Fig. 11c, d), altered SM cell alignment (Supplementary Fig. 11e, f) and shape (Supplementary Fig. 11e, g) as well as decreased F-actin levels (Supplementary Fig. 11e, h) compared to controls. We then used ouabain, a well established inhibitor of Na+/K+-ATPase^32^, to deplete intracellular potassium in the presence of elevated extracellular potassium. Notably, ouabain treatment, compared to DMSO treatment, partially rescued the SM cell alignment (Supplementary Fig. 11i, j) and shape (Supplementary Fig. 11i, k) phenotypes caused by the increase of extracellular potassium. These results further support the model that intracellular ion homeostasis is crucial for SM cell alignment and shape.

Smooth muscle contraction has been reported to drive tubulogenesis^35^. Based on the findings that *Kcnj13* is required for SM cell alignment and actin organization, we hypothesized that a disruption of SM cell orientation might lead to compromised circumferential tracheal contraction resulting in tube elongation defects in *Kcnj13^T38C/T38C^* mice. To examine SM contractility, we measured contractile forces in several settings. *Kcnj13^T38C/T38C^* tracheas exhibited greatly reduced contractile forces compared to WT (Supplementary Fig. 12a). Notably, after 50 μM VU590 treatment, WT tracheas also exhibited impaired SM contraction (Supplementary Fig. 12a). In addition, at early embryonic stages, the anterior part of *Kcnj13^T38C/T38C^* tracheas exhibited an expanded tube diameter compared to WT (Supplementary Fig. 12b), while the posterior part appeared slightly narrowed (Fig. 4d). Altogether, these data indicate that SM-mediated circumferential contraction may prevent tracheal tube over-expansion and promote its elongation along the longitudinal direction.

### p-AKT as a mediator of *Kcnj13* function in SM cells

We aimed to further understand how KCNJ13 could influence actin organization in tracheal SM cells. The Ser/Thr kinase AKT, an essential actin organizer^10,11^, has been reported to depend on potassium homeostasis for its activation state, as assessed by Ser473 phosphorylation^32,36^. We thus examined the phosphorylation level of AKT at Ser473 and found that it was greatly reduced in *Kcnj13^T38C/T38C^* tracheas compared to WT (Fig. 6a-d). In addition, after 50 μM VU590 treatment, WT tracheas also exhibited greatly reduced phosphorylation levels of AKT at Ser473 compared to controls (Fig. 6e, f). Furthermore, *Kcnj13^T38C/T38C^* tracheas, after treatment with A-443654, a small molecule that increases AKT phosphorylation at Ser473^37^, exhibited partially rescued SM cell phenotypes (Fig. 6g-i) and F-actin levels (Fig. 6j, k) compared to controls. These results indicate that reduced AKT phosphorylation in *Kcnj13^T38C/T38C^* tracheas partially accounts for the SM cell alignment and shape phenotypes via its action on actin polymerization (Fig. 6l).

**Figure 6.**
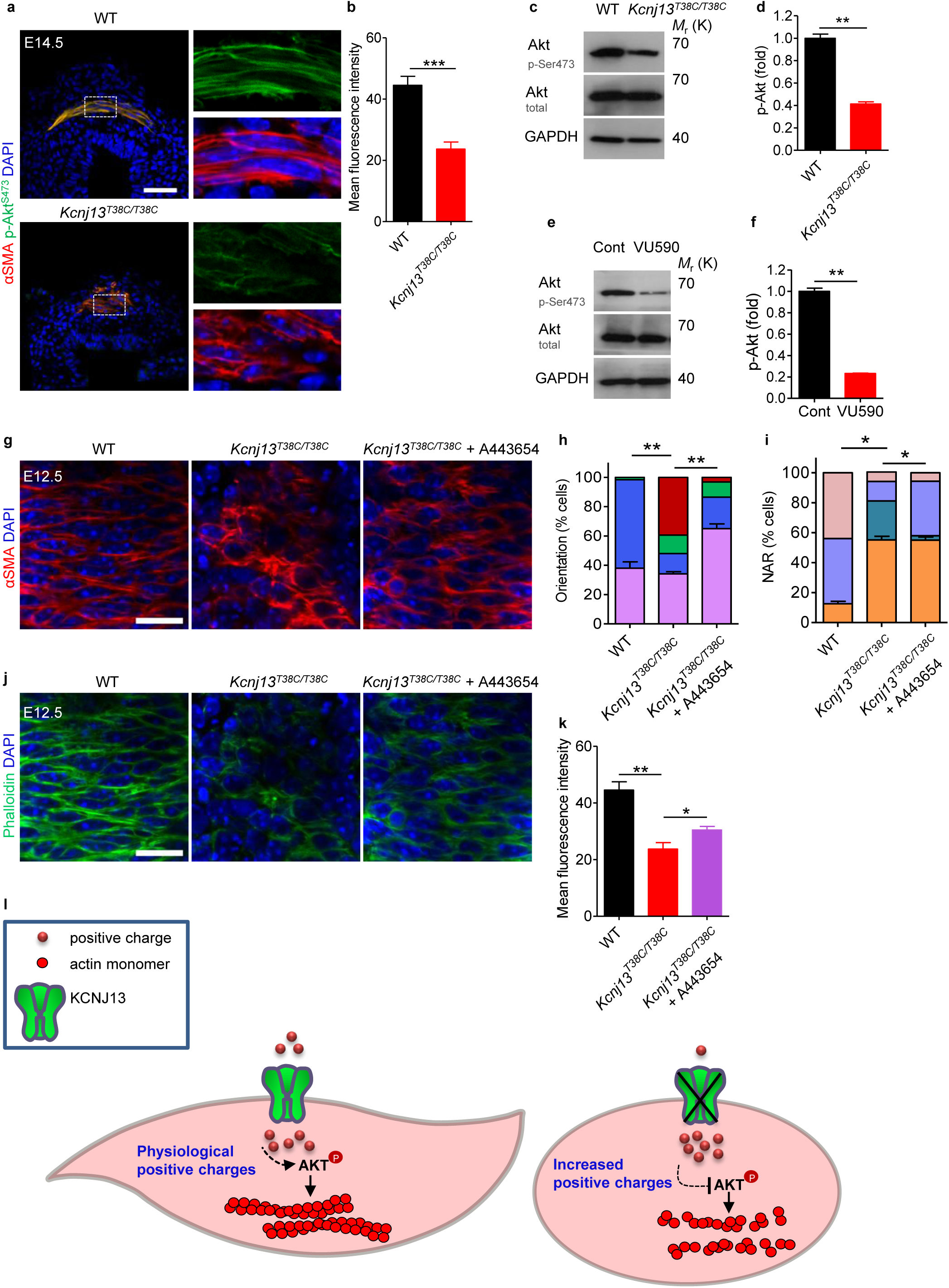
AKT phosphorylation as a mediator of *Kcnj13* function in tracheal SM cells. (**a**) Immunostaining for αSMA (red) and p-AKT^ser473^ (green) and DAPI staining (blue) of transverse sections of E14.5 WT (n=6) and *Kcnj13^T38C/T38C^* (n=6) tracheas. (**b**) Quantification of mean p-AKT^ser473^ fluorescence intensity in E14.5 WT (n=6) and *Kcnj13^T38C/T38C^* (n=6) tracheal SM cells. (**c**) Western blotting for phosphorylated AKT, total AKT and GAPDH in P0 WT (n=12) and *Kcnj13^T38C/T38C^* (n=12) tracheas. (**d**) Quantification of relative p-AKT^ser473^ levels in P0 WT (n=12) and *Kcnj13^T38C/T38C^* (n=12) tracheas. (**e**) Western blotting for phosphorylated AKT, total AKT and GAPDH in P0 tracheas after DMSO (n=16) or 50 μM VU590 (n=16) treatment. (**f**) Quantification of relative p-AKT^ser473^ levels in P0 tracheas after DMSO (n=16) or 50 μM VU590 (n=16) treatment. (**g**) Immunostaining for αSMA (red) and DAPI staining (blue) in dorsal views of E12.5 WT (n=16) and *Kcnj13^T38C/T38C^* (n=8) tracheas after a 48 hour DMSO treatment, and *Kcnj13^T38C/T38C^* (n=8) tracheas after a 48 hour 0.5 μM A443654 treatment. Quantification of SM cell orientation (**h**) and NAR (**i**) of E12.5 WT (n=8) and *Kcnj13^T38C/T38C^* (n=8) tracheas after a 48 hour DMSO treatment, and *Kcnj13^T38C/T38C^* (n=8) tracheas after a 48 hour 0.5 μM A443654 treatment. (**j**) Phalloidin (green) and DAPI staining (blue) in dorsal views of E12.5 WT (n=8) and *Kcnj13^T38C/T38C^* (n=8) tracheal SM cells after a 48 hour DMSO treatment, and *Kcnj13^T38C/T38C^* (n=8) tracheal SM cells after a 48 hour 0.5 μM A443654 treatment. (**k**) Quantification of mean phalloidin fluorescence intensity in SM cells (as in **j**). (**l**) A working model for the KCNJ13/AKT pathway in tracheal SM cells. KCNJ13 is expressed in tracheal SM cells to maintain intracellular positive charges at physiological levels, which leads to AKT phosphorylation and actin polymerization, which are essential for SM cell alignment and polarity. When KCNJ13 is inactivated, intracellular positive charge levels increase, leading to AKT hypophosphorylation and actin depolymerization, ultimately resulting in altered SM cell alignment and polarity during tubulogenesis. Scale bars: 50 μm (**a**), 20 μm (**g, j**). \**P* < 0.05; \*\**P* < 0.01; \*\*\**P* < 0.001; Unpaired Student’s *t*-test, mean ± s.d.

## Discussion

Understanding the mechanisms underlying tube formation is a fundamental goal in developmental biology. The reported roles of SM cells in tubulogenesis include wrapping nascent epithelial buds for terminal bifurcation during lung branching morphogenesis^38^, and folding the epithelium into villi via mucosal buckling during the formation of the gut tube^39^. *Kcnj13^T38C/T38C^* mice exhibit a shorter trachea but no obvious defects in cell division plane or cell proliferation in the tracheal epithelium, suggesting that loss of *Kcnj13* function affects some other cell type(s) during tracheal tubulogenesis. Indeed, we found that SM cell-specific inactivation of *Kcnj13* phenocopied some of the defects observed in the global mutants including a shorter trachea and altered SM alignment. Mutant animals also exhibit reduced F-actin levels in their tracheal SM cells as well as compromised circumferential contraction indicating major defects in these cells. However, it is important to note that the trachea of SM-specific *Kcnj13* mutants is longer than that of global mutants, suggesting that *Kcnj13* regulates additional cell type(s) necessary for tracheal elongation. Altogether, our data support a model whereby SM morphogenesis and contractility drive the elongation of the developing trachea; in *Kcnj13* mutants, as many SM cells lose their circumferential alignment, the elongation of the tube becomes compromised, possibly as a result of both reduced circumferential contraction and increased longitudinal alignment of the SM cells, the latter of which would block elongation, as in the gut tube^39^. Interestingly, *Wnt5a* and *Ror2* mutants also exhibit a shorter trachea^40,41^, a phenotype recently shown to be caused by defects in SM cell morphology and polarity^13^. It will thus be interesting to investigate whether and how *Wnt5a*/*Ror2* and *Kcnj13* intersect in the regulation of tracheal tubulogenesis including elongation. Additional studies have shown that deletion of TMEM16A, a calcium-activated chloride channel, leads to apparently shorter tracheas as well as altered tracheal SM cell shape and organization^42^, while CFTR, a chloride ion channel, leads to a reduced tracheal SM cell mass^43^ as well as fracture of the tracheal cartilage rings^44^, phenotypes all observed in *Kcnj13^T38C/T38C^* tracheas. It thus appears that multiple ion channels regulate tracheal SM cells and cartilage formation.

Our data on actin organization in tracheal SM cells suggest a model by which ion channels mediate actin filament reorganization through the actin organizer AKT. Although the exact mechanisms by which KCNJ13 regulates AKT activity remain to be determined, it is possible that KCNJ13 inactivation inhibits PI3K or activates PTEN, thereby impairing AKT plasma membrane translocation and subsequent activation. Other possibilities are that KCNJ13 inactivation leads to inhibition of the mTORC2 kinase^45^, or activation of the PHLPP phosphatase^46^.

In addition, point mutations in *KCNJ13* are associated with several human disorders^47–49^. Our *Kcnj13* point mutant (c. 38T>C (p.Leu13Pro)) exhibits severe primary tracheomalacia, providing a new model to investigate the etiology of this disease, and develop therapeutic approaches.

Overall, our data reveal the importance of potassium channels in regulating AKT phosphorylation, actin organization and epithelial tube formation, providing further insights into the relationship between ion homeostasis and cytoskeletal organization.

## METHODS

### Experimental animals

All mouse husbandry was performed under standard conditions in accordance with institutional (MPG) and national ethical and animal welfare guidelines. 30 C57BL/6J male mice treated with a 3 X 100 mg/kg dose of ENU^50^ were obtained from Dr. Monica Justice (Baylor College of Medicine, Houston, TX). After a period of 10 weeks for the recovery of fertility, the mutagenized G0 males were crossed with C57BL/6J female mice. G1 males were outcrossed with C57BL/6J females to generate G2 females. Four G2 females were backcrossed with their G1 father, and the resulting G3 P0 pups were subject to tracheal and lung dissection and analysis. *Nkx2.1^Cre^*, *Myh11-CreER^T2^* and *Shh^flox^* alleles have been previously described^6,21^. The *Kcnj13* deletion (*Kcnj13^Del-E2&3^*) and floxed (*Kcnj13^flox^*) alleles were generated using the CRISPR-Cas9 system and homology directed repair. Two gRNAs (targets #1 and #2) (Supplementary Fig. 2) were selected for mouse *Kcnj13* to direct Cas9 cleavage and insertion of loxP sites flanking exons 2 and 3 using an online CRISPR design tool (http://crispr.mit.edu/). The gRNA sequences were cloned into pDR274 (Addgene, Cambridge, MA) for gRNA production^51^. Next, gRNAs and Cas9 mRNA were synthesized and microinjected into the cytoplasm of C57BL/6 inbred zygotes^52^, together with two 110-bp single-stranded donors (#1 and #2). After injection, surviving zygotes were immediately transferred into oviducts of ICR albino pseudopregnant females. *Kcnj13^Del-E2&3^* and *Kcnj13^flox^* alleles were detected in G0 mice and germline-transmitted. For genotyping of *Kcnj13^DeL-E2&3^* mice, primers Fwd (5’-CATAAGAGTCAGCGCCTTCA-3’) and Rev (5’-AGGCTCAGCTAACCAAGCATGA-3’) were used to generate a ~350 bp PCR amplicon from the deletion allele. For genotyping of *Kcnj13* floxed mice, primers sets (loxP1-Fwd: 5’-AAAATTTTACTTCTCTCAACTTCT-3’; LoxP1-Rev: 5’-AAACATTTTTGGTTTTGTTTT-3’ and loxP2-Fwd: 5’-CAACTTAGATTTATGCTTGAAA-3’; LoxP2-Rev: 5’-AAATAGACATTGATGATGTTGTT-3’) were used to generate ~85 bp wild-type and ~125 bp loxP PCR amplicons for both sites. All breeding colonies were maintained under cycles of 12-hour light and 12-hour dark. All animal procedures for generating the *Kcnj13^DeL-E2&3^* and *Kcnj13^flox^* alleles were approved by the Institutional Animal Care and Use Committee at Baylor College of Medicine and all animal experiments were done in compliance with ethical guidelines and approved protocols.

### Whole-exome sequencing analysis

Genomic DNA from two WT and two mutant mice were isolated using a standard protocol, captured using Agilent SureSelect Mouse All Exon kit V1, and sequenced using Illumina HiSeq 2000 with minimum average 50× target sequence coverage (BGI-Hong Kong). Sequence reads were aligned to the C57BL/6J mouse reference genome (mm10) and analysed using CLCBio Genomic Workbench and GATK software. To minimize false negatives, variant calls were set at 5× minimum coverage and ≥20% alternate reads. Sequence variants were annotated to SNPs from dbSNP (version 142) and filtered against dbSNP128.

### Alcian blue staining of cartilage

Trachea cryosections (10 µm) were fixed in 4% paraformaldehyde for 20 minutes, treated with 3% acetic acid solution for 3 minutes, stained in 0.05% alcian blue for 10 minutes and counterstained with 0.1% nuclear fast red solution for 5 minutes. For whole-mount staining of tracheal cartilage, dissected tracheas were fixed in 95% ethanol for 12 hours followed by overnight staining with 0.03% alcian blue dissolved in 80% ethanol and 20% acetic acid. Samples were cleared in 2% KOH.

### Tracheal tube length measurements

Tracheal tube length was calculated by measuring the distance between the first and last tracheal cartilage rings.

### Whole-mount immunostaining

Tracheas were isolated from E11.5 to E16.5 embryos and P0 pups and fixed in DMSO:Methanol (1:4) overnight at 4°C. Whole tracheas were then incubated in H_2_O_2_/DMSO/methanol (1:1:4) for 5 hours at RT. After bleaching, samples were washed twice in 100% methanol for 1 hour each, once in 80% methanol for 1 hour, once in 50% methanol for 1 hour, twice in PBS for 1 hour each, and twice in 5%FBS/ PBS/0.5% Triton X-100/3%BSA for 1 hour each. To perform whole-mount immunostaining^53^, tracheas were incubated with primary antibodies diluted in 5% FBS/PBS/0.5% Triton X-100/3% BSA for 24 hours at 4 °C, washed five times for 1 hour each at 4°C with 5% FBS/PBS/0.5% Triton X-100/3% BSA followed by incubation with secondary antibodies for 24 hours at 4 °C, washed five times for 1 hour each at 4°C with 5% FBS/PBS/0.5% Triton X-100/3% BSA, dehydrated in methanol and then cleared in benzyl alcohol:benzyl benzoate (1:2). To visualize smooth muscle (SM) cells and tracheal mesenchymal cells, tracheas were stained for αSMA and SOX9, respectively. To visualize the Golgi apparatus, tracheas were stained for GM130. To visualize F-actin, tracheas were stained with 488-conjugated phalloidin (Thermo Fisher Scientific, A12379).

### Immunostaining of cryosections

Tracheas were dissected in PBS, fixed in 4% paraformaldehyde overnight at 4 °C, incubated in 10% sucrose and 30% sucrose for 24 hours each at 4 °C, mounted in OCT embedding compound, and sectioned at 10 µm. To perform immunostaining, sections were fixed in 4% paraformaldehyde 10 minutes at 4 °C, followed by incubation in permeabilization solution (0.3% Triton X-100/PBS) for 15 minutes at RT, incubated in blocking solution (5% FBS/PBS/3% BSA) for 1 hour at RT, incubated in primary antibodies overnight at 4 °C, washed, incubated in secondary antibodies for 2 hours at RT, washed, and then mounted for imaging. Immunostaining for p-AKT^Ser473^ and αSMA was carried out with WT and mutant trachea sections on the same slide.

### Airspace measurements

6 random fields per lung tissue section stained with hematoxylin and eosin from P0 WT and mutants were acquired with Axio Imager Z2. The percentage of terminal airspaces in the lung is the proportion of blank area of each field relative to the total area^13^.

### *In situ* mRNA hybridization of cryosections

Lungs were dissected in PBS, fixed in 4% paraformaldehyde overnight at 4 °C, mounted in OCT embedding compound, and then sectioned at 10 µm. To perform *in situ* hybridization^54^, cryosections were permeabilized in 5 μg/ml proteinase K (Roche) for 15 min at RT, followed by acetylation for 2 min and pre-incubation in hybridization buffer for 3 h at 70 °C, incubated with DIG-labeled RNA antisense probes overnight at 70 °C, washed, incubated with alkaline phosphatase-conjugated anti-digoxigenin antibody (Roche) overnight at 4°C, washed, and then the signal was detected with NBT-BCIP staining solution (Roche).

### Reverse transcription quantitative PCR (RT–qPCR)

Total RNA extraction was conducted using a miRNeasy Mini Kit (Qiagen). cDNA was synthesized using the Maxima First Strand cDNA Synthesis Kit (Thermo Fisher Scientific), according to manufacturer’s instructions. Quantitative real-time PCR was performed using Eco Real-Time PCR System (Illumina) and Maxima SYBR Green/Fluorescein qPCR Master Mix (Thermo Fisher Scientific). The following primers were used: *Actb* forward 5’-CGGCCAGGTCATCACTATTGGCAAC-3’ and *Actb* reverse 5’-GCCACAGGATTCCATACCCAAGAAG-3’; *Kcnj13* forward 5’-TGGTGAACTTTACCAGACCAGT-3’ and *Kcnj13* reverse 5’-GGATGTCCTCCTTTGGCAGAT-3’; *Wnt5a* forward 5’-CACTTAGGGGTTGTTCTCTGA-3’ and *Wnt5a* reverse 5’-ATATCAGGCACCATTAAACCA-3’; *Ror2* forward 5’-CCCAACTTCTACCCAGTCCA-3’ and *Ror2* reverse 5’-TGTCCGCCACAGATGTATTG-3’; *Fzd4* forward 5’- TTGTGCTATGTTGGGAACCA-3’ and *Fzd4* reverse 5’-GACCCCGATCTTGACCATTA-3’; *Fzd6* forward 5’-GTGCTGCAAGAGTCCTGTGA-3’ and *Fzd6* reverse 5’-CGCTGCTCTTTGGACTTACC-3’; *Fzd8* forward 5’-ACTACAACCGCACCGACCT-3’ and *Fzd8* reverse 5’-ACAGGCGGAGAGGAATATGA-3’; *Vangl1* forward 5’- GATGCTGTTAGGAGGTTCGG-3’ and *Vangl* reverse 5’-AGTCCCGCTTCTACAGCTTG-3’; *Dvl1* forward 5’-CCTTCCATCCAAATGTTGC-3’ and *Dvl1* reverse 5’-GTGACTGACCATAGACTCTGTGC-3’; *Dvl2* forward 5’-ACTGTGCGGTCTAGGTTTTGAGTC-3’ and *Dvl2* reverse 5’-GGAAGACGTGCCCAAGGA-3’; *Dvl3* forward 5’-AGGGCCCCTGTCCAGCT-3’ and *Dvl3* reverse 5’-AAAAGGCCGACTGATGGAGAT-3’; *Celsr1* forward 5’-ATGCTGTTGGTCAGCATGTC-3’ and *Celsr1* reverse 5’-GGGATCTGGACAACAACCG-3’; *Celsr2* forward 5’-GCTGTGTGTGAGCATCTCGT-3’ and *Celsr2* reverse 5’-CATCATGAGTGTGCTGGTGT-3’; *Pk1* forward 5’-GATGGAGAAAGCAAGCCAAG-3’ and *Pk1* reverse 5’-TGTGCAGCATGGAAGAGTTC-3’; *Fgf10* forward 5’- CGGGACCAAGAATGAAGACT-3’ and *Fgf10* reverse 5’-AGTTGCTGTTGATGGCTTTG-3’; *Hgf* forward 5’- AACAGGGGCTTTACGTTCACT-3’ and *Hgf* reverse 5’-CGTCCCTTTATAGCTGCCTCC-3’; *Pdgfa* forward 5’-GAGGAAGCCGAGATACCCC-3’ and *Pdgfa* reverse 5’-TGCTGTGGATCTGACTTCGAG-3’; *Tgfb1* forward 5’-GAGCCCGAAGCGGACTACTA-3’ and *Tgfb1* reverse 5’-TGGTTTTCTCATAGATGGCGTTG-3’.

### Western blotting

Isolated P0 tracheas were lysed using RIPA buffer (Cell Signaling, 9806) supplemented with protease and phosphatase inhibitors (Cell Signaling, 5872). Lysates were centrifuged at 10,000 g for 10 minutes, subjected to SDS-PAGE and transferred to nitrocellulose membranes. Membranes were probed with primary and HRP-conjugated secondary antibodies (Cell Signaling Technology) and were developed using the ECL detection system (Pierce). For chemical treatment, tracheas were incubated in DMEM/F-12 medium containing 0.1% DMSO or 50 μM VU590 for 4 hours before lysis. All uncropped images related to western blotting data are available in Supplementary Fig. 13.

### Quantification of western blot signals

The AKT, p-AKT and GAPDH levels were quantified using ImageJ. AKT and p-AKT levels were normalized to the values yielded by GAPDH. p-AKT fold change was calculated by the ratio of p-AKT/AKT and WT was assigned to 1.

### Antibodies

The following antibodies were used: Mouse anti-αSMA-Cy3 (1:1000, Sigma-Aldrich, C6198); Rat anti-CDH1 (1:500, Santa Cruz, sc-59778); Goat anti-KCNJ13 (1:50, Santa Cruz, C19); Rabbit anti-SOX9 (1:400, Millipore, AB5535); Sheep anti-GM130 (1:50, R&D systems, AF8199); Rabbit anti-Ki67 (1:400, Cell Signaling Technologies, #9027); Rabbit anti Cleaved Caspase-3 (1:600, Cell Signaling Technologies, #9661); Rabbit anti KRT5 (1:1000, Abcam, ab53121); Goat anti-CC10 (1:200, Santa Cruz, T-18); Rabbit anti-NKX2.1 (1:400, Santa Cruz, H-190); Rabbit anti-SFTPC (1:400, Millipore, AB3786); Mouse anti-PCNA (1:400, Santa Cruz, sc-56); Rabbit anti-PH3 (1:400, Millipore, 06-570); Rabbit anti-p-AKT^ser473^ (1:200, Cell Signaling Technologies, #4060); Rabbit anti-AKT (1:2000 Cell Signaling Technologies, #9272) and Rabbit anti-GAPDH (1:3000, Cell Signaling Technologies, #2118).

### Explant culture of mouse embryonic tracheas and lungs, and chemical treatment

Tracheas and lungs were isolated from E12.5 embryos and cultured using an established protocol^33^. For KCl (P9541, Sigma) treatment, a 2 M stock solution was diluted to 40 mM. For VU590 (3891, TOCRIS) treatment, a 40 mM stock solution was diluted to 40 μM. For valinomycin (3373, TOCRIS) treatment, a 2 mM stock solution was diluted to 2 μM. For A443654 (16499, Cayman Chemical) treatment, a 0.5 mM stock solution was diluted to 0.5 μM. Isolated tracheas and lungs were cultured in DMEM/F-12 medium containing the above chemicals at 37 °C in a 5% CO2 incubator for 48 hours. For ouabain (1076, TOCRIS) treatment, a 100 mM stock solution was diluted to 200 μM in DMEM/F-12 medium and isolated tracheas and lungs were pre-incubated for 30 minutes and then cultured for another 48 hours after adding 40 mM KCl. 0.1% DMSO in DMEM/F-12 medium was used as a control for VU590, valinomycin and A443654 treatment. 0.2% DMSO in DMEM/F-12 medium was used as a control for ouabain treatment. The medium was replaced every 24 hours before collection for analysis.

### Tracheal SM cell membrane potential measurements

For the examination of tracheal SM cell membrane potential, E14.5 tracheas and lungs were incubated in 10 μg/ml bis-(1,3-dibutylbarbituric acid) trimethine oxonol [DiBAC4(3)] (D8189, Sigma) diluted in PBS, in the dark for 30 minutes at RT. For VU590 treatment, tracheas and lungs were pre-incubated in 50 μM VU590 diluted in PBS for 2.5 hours and incubated for another 0.5 hour after adding 10 μg/ml DiBAC4(3). For KCl treatment, tracheas and lungs were pre-incubated in 40 mM KCl diluted in PBS for 1 hour and incubated for another 0.5 hour after adding 10 μg/ml DiBAC4(3). Samples were washed twice in PBS for 10 minutes each before collection for analysis.

### Imaging

Imaging of wholemount tracheas and trachea sections was performed using a Nikon SMZ25 or Zeiss 880 upright laser scanning confocal microscope. Quantification of lumen area, tube length, SM area, SM cell orientation and nuclear aspect ratio (NAR), Golgi-apparatus position relative to the nucleus, and immunofluorescence intensity was performed using ImageJ (http://rsbweb.nih.gov/ij/).

### *Ex vivo* trachea physiology

2 mm sections of tracheas were isolated from P0 pups and kept in Krebs solution (119 mM NaCl, 4.7 mM KCl, 2.5 mM CaCl_2_, 1.17 mM MgSO_4_, 20 mM NaHCO_3_, 1.18 mM KH_2_PO_4_, 0.027 mM EDTA, 11 mM glucose) aerated with carbogen at 37 °C. Tracheal rings were mounted in a wire-myograph system (610-M, Danish Myo Technology) and a resting tension of 2 mN was applied for each ring as a baseline. Contractile responses were determined by cumulative administration of indicated acetylcholine concentrations. For studies where VU590 was used, tracheas were incubated with 50 μM VU590 for 10 minutes before acetylcholine exposure.

### Sample size

Sample sizes for animal studies were made as large as possible based on the complex genetics. Experiments in the manuscript have n≥5 to ensure robustness. For all animal experiments, multiple animals were tested in 2–4 individual experiments.

### Sample or animal exclusions

No samples or animals were excluded from the analysis.

### Randomization

For all animal experiments, WT and mutant animals were chosen at random from among littermates with the appropriate genotype.

### Blinding

The genotype of WT and mutant mice was unknown to the investigators before data analysis in each experiment. In the KCl and VU590 treatment experiments, control and KCl or VU590 treated mice were littermates with the same genotype.

### Statistical analysis

No statistical methods were used to predetermine sample size. Statistical analyses were performed using GraphPad software. Error bars, s.d. P values were calculated by two-tailed Student’s *t*-test (*P<0.05; **P<0.01; ***P<0.001; ****P<0.0001; NS, not significant). P<0.05 indicates a finding is significant. P>0.05 indicates a finding is not significant.

### Data Availability

The authors declare that all data supporting the findings of this study are available within the article and its supplementary information files or from the corresponding author upon reasonable request.

## ACKNOWLEDGEMENTS

We thank Brigid Hogan, Jason Rock, Xin Sun, Pao-Tien Chuang, Ross Metzger, Cecilia Lo, Kathryn Anderson, Diana Laird, Paul Potter, Monica Justice and Joaquim Grego-Bessa for discussion and advice on the lung ENU screen, Radhan Ramadass, Jenny Pestel and Yu Hsuan Carol Yang for imaging assistance, Thomas Braun for reagents, Rubén Marín-Juez, Zacharias Kontarakis, Andrea Rossi, Jason Kuan Han Lai, Paolo Panza, Konstantinos Gkatzis, Sabine Fisher, Sharon Meaney-Gardian and Nana Fukuda for discussion and/or assistance. Funding for this study was provided in part by the Max Planck Society to S.O. and D.Y.R.S.

## AUTHOR CONTRIBUTIONS

W.Y. and D.Y.R.S. conceived the project, designed experiments and analyzed data; W.Y., H.-T.K., F.G., W.L., B.G., C.B., V.G., G.S., S.G. and S.O. contributed to experiments and data analysis; K.K. and M.M. shared their data regarding SM cell polarity and tracheal elongation prior to publication; S.W. performed western blot and trachea physiology experiments and data analysis; H.Z., D.R. and G.M. generated and characterized *Kcnj13^flox^* and *Kcnj13^Del-E2&3^* mice, and contributed to data analysis; J.P. and M.L. performed data analysis for whole-exome sequencing; W.Y. and D.Y.R.S. wrote the manuscript. All authors commented on the manuscript.

